# Spatial Analysis of Functional Enrichment (SAFE) in Large Biological Networks

**DOI:** 10.1101/094904

**Authors:** Anastasia Baryshnikova

## Abstract

Spatial Analysis of Functional Enrichment (SAFE) is a systematic quantitative approach for annotating large biological networks. SAFE detects network regions that are statistically overrepresented for functional groups or quantitative phenotypes of interest, and provides an intuitive visual representation of their relative positioning within the network. In doing so, SAFE determines which functions are represented in a network, which parts of the network they are associated with and how they are potentially related to one another.

Here, I provide a detailed stepwise description of how to perform a SAFE analysis. As an example, I use SAFE to annotate the genome-scale genetic interaction similarity network from *Saccharomyces cerevisiae* with Gene Ontology (GO) biological process terms. In addition, I show how integrating GO with chemical genomic data in SAFE can recapitulate known modes-of-action of chemical compounds and potentially identify novel drug mechanisms.

## Introduction

Network diagrams have long been used to represent molecular structures, signaling cascades, biochemical reactions and many other types of connections between components of biological systems. In all of their variations, ball-and-stick models had unparalleled success in illustrating complex relationships and revealing their organizational structure. With the rise of genomic technologies, biological networks grew exponentially in size and density, but our understanding of their global organization did not improve at the same rate. One likely reason is our inability to read and interpret large networks in the same way we do it for small ones, i.e. by locating familiar nodes and inferring their relationship to unfamiliar ones based on distance and connectivity. Using the same approach with genome-scale networks is highly impractical due to the mere number of nodes, the intricacy of their connections and the inconsistency of our knowledge about biological functions.

To address this problem and expand our understanding of complex biological networks, I have recently developed a strategy named Spatial Analysis of Functional Enrichment (SAFE) (Baryshnikova, 2016). This method is designed to provide a birdseye view of the functional organization of a network by mapping the distribution of biological knowledge throughout all of its local neighborhoods. In doing so, SAFE relies on three basic ideas: 1) that nodes in a large network can be spatially organized in a way that reflects their connectivity; 2) that the resulting node positions can be used to identify network neighborhoods associated with prior knowledge; 3) that these associations can be visually represented on the network and drive its biological interpretation. In practice, SAFE detects network regions that are statistically overrepresented for a functional group or a quantitative phenotype of interest, and provides an intuitive visual representation of their relative positioning within the network.

In a proof-of-concept experiment, I used SAFE to systematically examine the genome-wide genetic interaction similarity (GIS) network from yeast *Saccharomyces cerevisiae* (Costanzo et al., 2010). By annotating the network with ∼4,400 Gene Ontology (GO) biological process terms, SAFE uncovered ∼20 large-scale domains associated with major cellular processes (Baryshnikova, 2016). SAFE also showed that different GO terms have different patterns of distribution throughout the GIS network and that these patterns can be potentially used to define term specificity (Baryshnikova, 2016). Furthermore, SAFE combined GO annotations with gene expression, chemical genomic and genetic interaction data to explore unfolded protein response (DeRossi et al., 2016) and identify a potentially novel cause of resistance to chemotherapy (Baryshnikova, 2016).

Here, I provide a practical demonstration of how to use SAFE for annotating a biological network. As an example, I use the most recent yeast GIS network (Costanzo et al., 2016) that is 40% larger and 50% denser than the one examined previously (Costanzo et al., 2010). In Example 1, I describe the steps required to reproduce the GO annotation of the new GIS network as shown in Figure 1F in (Costanzo et al., 2016). In Example 2, I illustrate how SAFE can integrate GO annotations with chemical genomic data to uncover drug mode-of-action.

**Figure 1.**
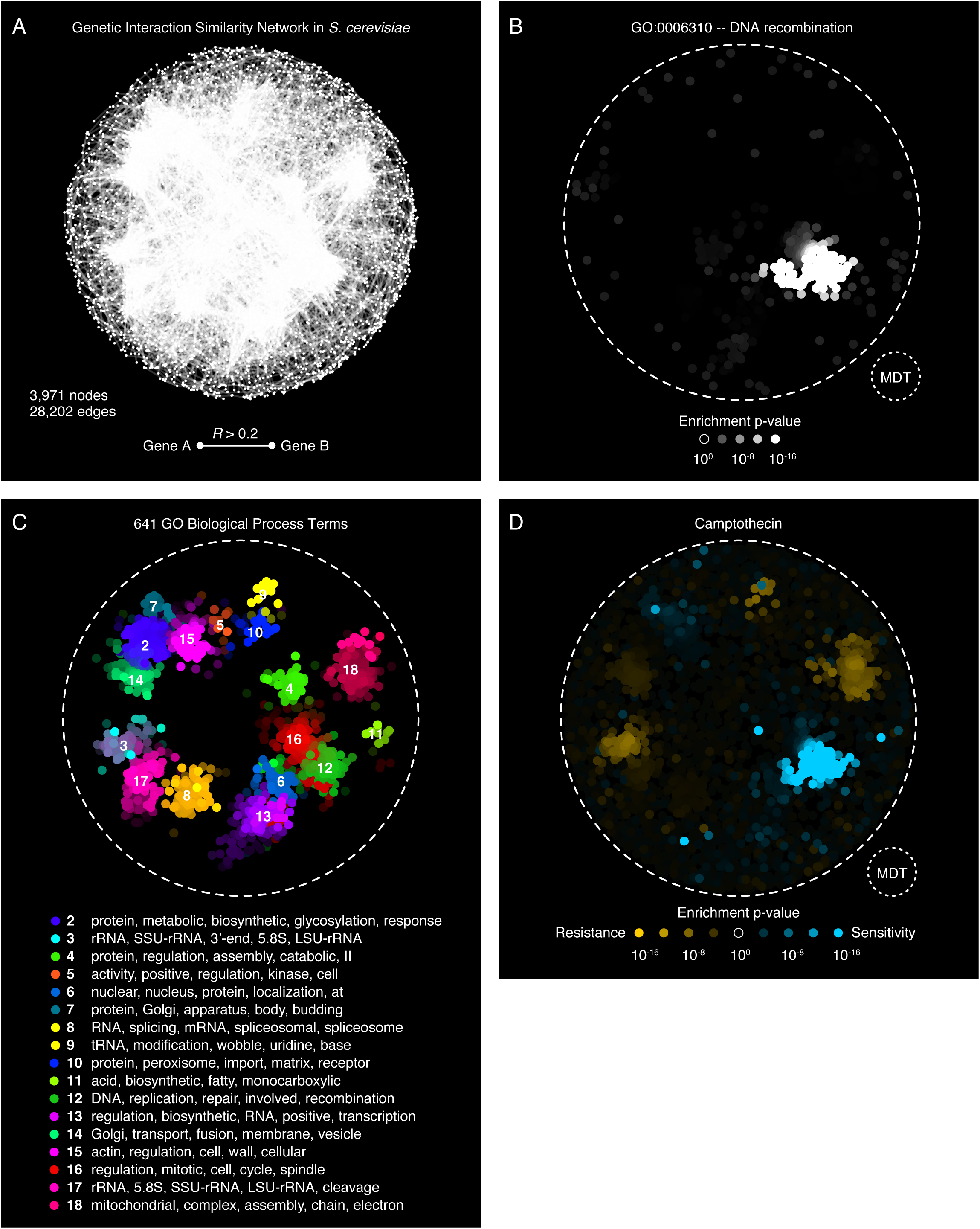
**(A)** The genetic interaction similarity (GIS) network, containing 3,971 nodes and 28,202 edges, was originally constructed in (Costanzo et al., 2016). Nodes represent genes. Two genes are connected if their genetic interaction profiles showed a Pearson correlation coefficient **(R)** greater than 0.2. The positions of the nodes in twodimensional space were determined by the spring-embedded layout algorithm, implemented in Cytoscape (Shannon et al., 2003). (B) An example of a network enrichment landscape produced by SAFE. The local neighborhood of every node in the network is tested for enrichment for the GO biological process term “DNA recombination” (GO:0006310). The resulting enrichment p-value is corrected for multiple testing, log-transformed into an enrichment score and assigned back to the node. The GO:0006310 enrichment scores of all nodes constitute the term’s enrichment landscape. In this example, the landscape is region-specific. All edges are hidden. All nodes are colored based on their enrichment scores. The small circle in the bottom right corner illustrates the approximate size of a neighborhood, defined by a set of nodes that are within a maximum distance threshold (MDT) from each other. **(C)** The final annotated network. SAFE combined 641 region-specific GO terms into 17 functional domains (labeled 2 to 18) based on the similarity of their enrichment landscapes. Different colors represent different domains. Each domain is labeled with a tag list, composed of the five words that occur most frequency within the names of the associated GO terms. The complete list of GO terms in each domain is provided in one of the output files. **(D)** The enrichment landscape of camptothecin, a chemical compound that inhibits DNA topoisomerase I. A chemical genomic screen measured the relative growth of ∼5,000 yeast knock-out mutants in the presence of camptothecin (Hoepfner et al., 2014). SAFE used the quantitative mutant growth data to annotate the GIS network. All edges are hidden. All nodes are colored based on the enrichment of their neighborhoods for genes that confer sensitivity (blue) or resistance (yellow) to camptothecin. The brightness of the color is proportional to the node’s enrichment score.

## Materials

### Software requirements

- SAFE: available as MATLAB code at https://bitbucket.org/abarysh/safe
- MATLAB R2014b or later
- Bioinformatics Toolbox for MATLAB
- Statistics and Machine Learning Toolbox for MATLAB
- Optional. Cytoscape 3.3 or later

### Input data files

- The Cytoscape session file containing the genetic interaction similarity network described in Costanzo et al., 2016 (Costanzo et al., 2016): included as part of the SAFE package (`/data/` folder) and available separately at https://bitbucket.org/abarysh/safe/master/data/Costanzo_Science_2016.cys
- Gene Ontology (GO) biological process annotations for yeast: included as part of the SAFE package (`/data/` folder) and available separately at https://bitbucket.org/abarysh/safe/master/data/go_bp_140819.mat
- The functional annotation file containing the chemical genomic screen for camptothecin described in Hoepfner et al., 2014 (Hoepfner et al., 2014): available at http://yeastphenome.org/datasets/1099/

## Methods

### SAFE installation, setup and basic workflow

1. Download the latest version of SAFE from Bitbucket and place it anywhere on your computer (e.g., in `/AAA/BBB/safe/`).
2. Open MATLAB and add the SAFE folder to the path: `addpath(genpath(‘/AAA/BBB/safe/’));`
3. Anywhere on your computer, create a folder that will store the results of the SAFE analysis: `mkdir(‘/CCC/DDD/’);`
4. Modify `/AAA/BBB/safe.m` (the main SAFE script) to record the absolute path to the results folder: `path_to_results_folder = ‘/CCC/DDD/’;`
5. Make a copy of the SAFE settings file (`/AAA/BBB/safe/safe.ini`) and save it in the results folder (`/CCC/DDD/safe.ini`). This file defines all tunable parameters and serves as a log record for every session of SAFE analysis. Do not change the name of the file. Also, do not remove the original file as it can be used as a reference for default parameters.
6. Modify `/CCC/DDD/safe.ini` in any text editor to adjust input/output parameters as needed (see Examples below for the most common settings).
7. Run `safe.m` (see also Note 2 and the examples below).
8. The results will be visually displayed in MATLAB and stored in `/CCC/DDD/`.

### Example 1. Annotating the GIS network with GO biological process terms

In this example, SAFE is used to annotate the yeast genetic interaction similarity (GIS) network with GO biological process terms. The steps of the analysis are run manually one-by-one to observe and explain the progression of the algorithm. The end result reproduces Figure 1F in (Costanzo et al., 2016).

1. Perform points 1–5 from the “SAFE installation and setup” section.
2. Make sure that the input network file (`Costanzo_Science_2016.cys`) is located at `/AAA/BBB/data/`. If not, see Materials to download the file. See also Note 1.
3. Modify `/CCC/DDD/safe.ini` to include the following changes (see below for a detailed explanation):
  a. `networkfile = /AAA/BBB/data/Costanzo_Science_2016.cys`
  b. `neighborhoodRadius = 7.5`
  c. `neighborhoodRadiusType = diameter`
  d. `unimodalityType = hartigans`
  e. `groupMinimize = 2`
4. Run Step 1 of `safe.m` (“Load settings”). See also Note 2.
  a. A subfolder with a unique SAFE session id is automatically created inside the results folder (e.g., `/CCC/DDD/safe-2016-11-02-16-08-08/`).
  b. All parameters from `safe.ini` are loaded into the main structure array (`layout`). This array will eventually contain all intermediate and final SAFE results, and will be automatically stored in `/CCC/DDD/<safe-session-id>/` together with other output files.
5. Run Step 2 of `safe.m` (“Load network”).
  a. The network is extracted from `networkfile` (specified in `safe.ini`) and loaded into `layout`.
  b. In this example, the genetic interaction similarity network containing 3,971 nodes and 28,202 undirected edges is loaded from the Cytoscape session file `Costanzo_Science_2016.cys`. A CYS file is effectively an archive of numerous XML and XGMML files. SAFE uncompresses the archive into the `/CCC/DDD/safe-session-id/` subfolder and parses the XGMML files containing information about nodes (labels and coordinates) and edges. See also Note 3.
  c. The following key fields in `layout` are populated:
    - `label`: the list of 3,971 node labels;
    - `label_orf`: the list of 3,971 systematic node labels that will be used to match nodes to functional annotations; systematic node labels do not have to be unique across the network;
    - `x` and `y`: the list of 3,971 node coordinates;
    - `edges`: a binary 3,971 × 3,971 network adjacency matrix.
6. Skip Step 3 of `safe.m` (“Apply network layout, if necessary”).
  a. In this example, the edge-weighted spring embedded layout has already been applied. See also Note 4.
7. Run Step 4 of `safe.m` (“Plot network”).
  a. A figure displays a black-and-white representation of the network (Figure 1A).
8. Skip Step 5 of `safe.m` (“Translate node labels, if necessary”).
  a. The list of systematic node labels (`label_orf`) has already been obtained from the Cytoscape session. See Note 5.
9. Run Step 6 of `safe.m` (“Load functional annotations”).
  a. The functional annotations are extracted from `annotationfile` (specified in `safe.ini`) and loaded into `layout`.
  b. In this example, `annotationfile` is empty, indicating that the default functional annotation standard should be loaded. The default standard corresponds to 4,373 GO biological process terms that were associated with at least 1 yeast gene as of August 19, 2014. When a gene is annotated to a GO term, it is automatically annotated to all of its parent terms. See also Note 6.
  c. The following key fields are populated in `layout`:
    - `group_ids`: the list of 4,373 GO term ids;
    - `group_names`: the list of 4,373 GO term names;
    - `label2group`: a binary 3,971 × 4,373 matrix of node-to-GO term annotations.
10. Run Step 7 of `safe.m` (“Calculate distances between node pairs”).
  a. Distance between all pairs of nodes is calculated based on the `nodeDistanceType` metric specified in `safe.ini`.
  b. In this example, `nodeDistanceType = shortpath_weighted_layout`. According to this metric, edges are weighted by their physical lengths resulting from the network layout, and the distance between two nodes is defined by the shortest physical path connecting them. See also Note 7.
  c. The following key field is populated in `layout`:
    - `nodeDistance`: a quantitative 3,971 × 3,971 matrix of node distances.
11. Run Step 8 of `safe.m` (“Calculate enrichments”).
  a. Every node in the network is associated with a local neighborhood, defined as the set of nodes located within a certain distance from it. The distance threshold is set by `neighborhoodRadiusType` and `neighborhoodRadius` in `safe.ini`. Each neighborhood is tested for enrichment of each attribute using a one-tailed Fisher’s exact test (see also Note 8), using the complete network as background (see Note 9). The corresponding p-values are corrected for multiple testing and log-transformed into enrichment scores. The set of enrichment scores for a given GO term across all neighborhoods constitutes the term’s enrichment landscape.
  b. In this example, `neighborhoodRadiusType = diameter` and `neighborhoodRadius = 7.5`. These settings indicate that two nodes are part of the same neighborhood if their distance is equal to or shorter than 7.5% of the network diameter. Network diameter is the distance between the two farthest nodes in the network. The threshold of 7.5% was defined arbitrarily; however, systematic evaluations showed that enrichment scores are robust to 2-fold variations of the distance threshold in either direction (Baryshnikova, 2016). See also Note 10.
  c. The following key fields are populated in `layout`:

- `R`: the distance threshold used to define neighborhoods;
- `neighborhoods`: a binary 3,971 × 3,971 matrix that defines neighborhood membership;
- `pval`: a quantitative 3,971 × 4,373 matrix of enrichment p-values;
- `opacity`: a quantitative 3,971 × 4,373 matrix of enrichment scores;
- `thresholdOpacity`: the lowest enrichment score to be considered significant (corresponds to p-value = 0.05);
- `opacity_01`: a binary 3,971 × 4,373 matrix of significant enrichments.
12. Run Step 9 of `safe.m` (“Plot sample enrichment landscapes”).
  a. A series of figures display the enrichment landscapes of several GO terms. These terms can be selected at random and/or specified by the user. The figures are exported to PDF and saved in the `/CCC/DDD/<safe-session-id>/` folder. The enrichment scores for all GO terms are exported into a tab-delimited text file and saved in the `/ CCC/DDD/<safe-session-id>/` folder as well.
  b. For example, the enrichment landscape of the GO term “DNA recombination” (GO:0006310, Figure 1B) can be plotted by executing the following command:
    - `plot_sample_groups(layout, ‘Range’, [find(layout.group_ids == 6310]);`
  c. Note that this step is optional. To suppress the figures when running `safe.m` as a script, set `plotSampleGroups` in `safe.ini` to `0`.
13. Run Step 10 of safe.m (“Combine functional attributes into domains”).
  a. First, the attributes are filtered based on the size and the pattern of their enrichment landscapes.
    - SAFE retains only attributes that are enriched in a minimum number of node neighborhoods (`groupEnrichmentMinSize` in `safe.ini)`. In this example, `groupEnrichmentMinSize = 10`.
    - If required (see Note 11), SAFE retains only attributes that are enriched in a single region of the network and excludes all attributes that are enriched in multiple regions or a broadly distributed. The method for measuring the enrichment distribution is set by `unimodalityType` in `safe.ini`. In this example, `unimodalityType = hartigans`.
  b. Second, SAFE constructs functional domains by grouping attributes with redundant enrichment landscapes, i.e. landscapes that are specific to the same region of the network. The grouping is done by computing pair-wise similarity between enrichment landscapes and performing hierarchical clustering. The similarity metric and the cluster threshold are set by `groupDistanceType` and `groupDistanceThreshold` in `safe.ini`, respectively. In this example, `groupDistanceType = jaccard` and `groupDistanceThreshold = 0.75` (see Note 12).
  c. Finally, functional domains are filtered by size and consolidated. The size of a functional domain is determined by how many node neighborhoods have their highest enrichment for an attribute assigned to that domain. Note that a node neighborhood can be enriched for multiple attributes in multiple domains; however, the attribute and the domain that show the highest enrichment will be considered predominant and officially “claim” the neighborhood. If a domain has claimed less than 10 node neighborhoods, it is excluded from further evaluation. After all the exclusions, the domains are re-numbered consecutively. See Note 13.
  d. The following key fields are populated in `layout`:
    - `grouplsTop`: a list of 4,373 values that indicate whether or not an attribute is included in the analysis (i.e., is region-specific and meets the minimum size criteria);
    - `groupColor`: a list of 4,373 domain ids, one for each attribute (“1” denotes attributes that were not assigned to any domain);
    - `labelColor`: a list of 3,971 predominant domain ids, one for each node (“1” denotes nodes that were not claimed by any domain);
    - `labelOpacity`: a list of 3,971 predominant enrichment scores, one for each node.
14. Run Step 11 of `safe.m` (“Generate automatic domain labels”).
  a. Each domain is provided with a summary label composed of the five most frequently occurring words within the labels of its attributes. Articles and other common words are excluded.
15. Run Step 12 of `safe.m` (“Plot annotated network”).
  a. A figure displays the final annotated network (Figure 1C). Nodes are positioned at their original x and y coordinates and colored by their average RGB value. For every domain, a numeric id is placed next to its center.
  b. To produce the figure, a random color is assigned to each domain and, by extension, to all of its associated attributes. Note that this random assignment results in a different (but equivalent) color palette at every run.
  c. For every node, attribute colors are combined proportionally to the enrichment score of each attribute in the node’s neighborhood. For example, if a node’s neighborhood is significantly enriched for attributes A, B and C, assigned to domains 3, 3 and 5, respectively, the color of the node will equal to (c_3_ * W_A_ + C_3_ * W_B_ + C_5_ * W_C_)/3, where c_3_ and c_5_ are the colors of domains 3 and 5 (in RGB format), and W_X_ is the square of the node’s enrichment score for attribute X.
16. Run Step 13 of `safe.m` (“Print results”).
  a. The annotated network is exported to PDF format and saved in `/CCC/DDD/<safe-session-id>/figure3.pdf`.
  b. Three additional output files are added to `/CCC/DDD/<safe-session-id>` (see also Note **Error! Reference source not found.**):
    - `domain_properties_annotation-highest.txt`: lists the properties of all functional domains (domain id, domain label and domain color, as it appears on the annotated network, in RGB format);
    - `attribute_properties_annotation-highest.txt`: lists the properties of all functional attributes (attribute id, attribute label and its corresponding domain id);
    - `node_properties_annotation-highest.txt`: lists the properties of all nodes (node unique label, node systematic label, id of predominant domain, enrichment score of predominant attribute, total number of associated domains and number of associated attributes per domain).
17. Run Step 14 of `safe.m` (“Save SAFE session”).
  a. This step saves the `layout` array into the `/CCC/DDD/<safe-session-id>/safe_session.mat` file.

### Example 2. Annotating the GIS network with camptothecin chemical genomic data

In this example, SAFE is used to annotate the yeast GIS network with a genome-wide chemical genomic screen for camptothecin, a DNA topoisomerase inhibitor that blocks DNA synthesis (Hoepfner et al., 2014). A chemical genomic experiment measures the relative growth of ∼5,000 yeast knock-out mutants in the presence of a chemical compound (Giaever et al., 2002). The sets of mutants that are relatively sensitive or resistant to the compound are often indicative on the compound’s mode-of-action and have been instrumental for identifying direct drug targets (Ho et al., 2011). Here, we annotate the GIS network with camptothecin growth data to identify network regions that are specifically enriched for sensitivity and resistance phenotypes. The steps of the analysis are run manually one-by-one to observe and explain the progression of the algorithm.

1. Repeat points 1–5 from the “SAFE installation and setup” section.
2. Make sure that the input network file (`Costanzo_Science_2016.cys`) is located at `/AAA/BBB/data/`. If not, see Materials to download the file. See also Note 1.
3. Modify `/CCC/DDD/safe.ini` to include the following changes (explained throughout the procedure):
  a. `networkfile = /AAA/BBB/data/Costanzo_Science_2016.cys`
  b. `annotationfile = /CCC/DDD/hoepfner_movva_2014_camptothecin.txt`
  c. `annotationsign = both`
  d. `unimodalityType = <empty>`
  e. `groupDistanceType = <empty>`
  f. `groupMinimize = 0`
4. Repeat points 4-8 from Example 1.
5. Run Step 6 of `safe.m` (“Load functional annotations”).
  a. The functional annotations are extracted from `annotationfile` (specified in `safe.ini`) and loaded into `layout`.
  b. In this example, `annotationfile` points to a file containing 1 functional attribute, i.e. the chemical genomic screen for camptothecin. The file lists 4,960 yeast knock-out mutants and their relative growth values in the presence of 200 uM camptothecin. Negative values indicate sensitivity, while positive values indicate resistance. Considered that we are interested in detecting network regions that are enriched for both phenotypes, we set `annotationsign = both`.
  c. The following key fields are populated in `layout`:
    - `group_ids`: 1 attribute id;
    - `group_names`: 1 attribute name.
    - `label2group`: a quantitative 3,971 × 1 matrix of node-to-attribute annotations.
6. Run Step 7 of `safe.m` (“Calculate distances between node pairs”).
  a. The following key field is populated in `layout`:
    - `nodeDistance`: a quantitative 3,971 × 3,971 matrix of node distances.
7. Run Step 8 of `safe.m` (“Calculate enrichments”).
  a. Every node in the network is associated with a local neighborhood, defined as the set of nodes located within a certain distance from it. The distance threshold is set by `neighborhoodRadiusType` and `neighborhoodRadius` in `safe.ini`. Within each neighborhood, the attribute values are summed to produce a quantitative neighborhood score. This score is compared to the distribution of 1,000 random scores obtained by randomizing node labels, and thus attribute values, across the entire network. The resulting empirical p-values are corrected for multiple testing and log-transformed into neighborhood enrichment scores. The set of enrichment scores for a given attribute across all neighborhoods constitutes the attribute’s enrichment landscape.
  b. In this example, `neighborhoodRadiusType = percentile` and `neighborhoodRadius = 0.5`. These settings indicate that two nodes are part of the same neighborhood if their distance is within the lowest 0.5th percentile of all pair-wise node distances. Also, given that `annotationsign = both` (see points 3 and 5 above), SAFE tests both tails of the random score distribution and produces 2 p-values for each neighborhood:
    - p-value #1 that measures the probability of observing a neighborhood score this low or lower in a random score distribution;
    - p-value #2 that measures the probability of observing a neighborhood score this high or higher in a random score distribution.
  c. The following key fields are populated in `layout`:
    - `R`: the distance threshold used to define neighborhoods;
    - `neighborhoods`: a binary 3,971 × 3,971 matrix that defines neighborhood membership;
    - `pval`: a quantitative 3,971 × 1 × 2 matrix of enrichment p-values (3,971 is the number of node neighborhoods, 1 is the number of attributes, 2 is the number of tails of the random distribution that were considered);
    - `opacity`: a quantitative 3,971 × 1 × 2 matrix of enrichment scores;
    - `thresholdOpacity`: the lowest enrichment score to be considered significant (corresponds to p-value = 0.05);
    - `opacity_01`: a binary 3,971 × 1 × 2 matrix of significant enrichments.
8. Run Step 9 of `safe.m` (“Plot sample enrichment landscapes”).
  a. A figure displays the enrichment landscape for camptothecin (Figure 1D). Given that `annotationsign = both` and each node has 2 enrichment scores (one for sensitivity and one for resistance), the landscape is produced by selecting the score with the highest value.
  b. The camptothecin enrichment landscape shows that the network region associated with DNA replication, recombination and repair (Figure 1C, domain 12), is statistically enriched for genes that confer sensitivity to the drug, when mutated (Figure 1D, blue). Similarly, network regions associated with ribosome biogenesis, tRNA modification and mitochondrial functions (Figure 1C, domains 3, 9 and 18) are enriched for resistance (Figure 1D, yellow).
  c. The figure is exported to PDF and saved in `/CCC/DDD/<safe-session-id>/figure2.pdf`.
  d. The enrichment scores are exported into two tab-delimited text files (one for the highest attribute values and one for the lowest) and saved in the `/CCC/DDD/<safe-session-id>/` folder as well.
9. Given than only 1 attribute is considered in this analysis, the remaining steps (“Combine functional attributes into domains” and others) can be skipped.

## Notes

1. In this example, we are using an existing Cytoscape session file in which the genetic interaction similarity network has been loaded and spatially organized using the edge-weighted spring embedded layout (Cytoscape.org, 2016), an implementation of the Kamada-Kawai algorithm (Kamada and Kawai, 1989). This Cytoscape session file was originally used for SAFE analysis in Costanzo et al., 2016 (Costanzo et al., 2016) and thus has the advantage of reproducing exactly the same outcome. As an alternative to importing the network from this file, the user can choose one of several options.
  a. Create a new Cytoscape session by loading the network from a different file format and applying the same or a different layout algorithm (see Cytoscape documentation for a list of available options (Shannon et al., 2003)).
  b. Load the network directly from a tab-delimited text file. SAFE accepts networks in list and adjacency matrix formats. In all cases, all edges with weights greater than 0 will be used.
    - 3-column list format: <Node 1 systematic label><Node 2 systematic label><Edge weight>
    - 5-column list format: <Node 1 label><Node 1 systematic label><Node 2 label><Node 2 systematic label><Edge weight>
    - Adjacency matrix: 1 row header (<Node 1 systematic label>), 1 column header (<Node 2 systematic label>), edges weights
2. The main SAFE file `/AAA/BBB/safe.m` can be executed as a regular MATLAB script (`run(‘/AAA/BBB/safe.m’);`). However, it is often more useful to manually run the different sections of the file, one step at a time, to get a better understanding of the algorithm and examine intermediate results.
3. The `Costanzo_Science_2016.cys` file contains only one network collection (`Costanzo_Science_2016_network_collection`) and only one network within that collection (`Costanzo_Science_2016_network`). However, in general, Cytoscape sessions can contain multiple network collections with multiple networks each. By default, SAFE will load the first network of the first network collection. To change the default, the user can modify `networkname` and `viewname` in `safe.ini` (see a more detailed explanation within `safe.ini` itself).
4. To reapply the layout and obtain a different configuration of the network, the user should set `layoutAlgorithm` in `safe.ini` to `Kamada-Kawai (Cytoscape`). Due to an initial randomization step, the Kamada-Kawai algorithm is non deterministic and results in a different network configuration at every run. To make layouts reproducible, the user should set `randomSeed` in `safe.ini` to any number (e.g., `randomSeed = 3`). This setting will make sure that, at every instance of SAFE, the nodes are initialized at the same random positions and will converge to the same final configuration.
5. Systematic node labels (`label_orf`) are used to match nodes to annotations (Step 6) and calculate enrichments (Step 8). As a result, it is important that `label_orf` is populated with the same label type that is used by the functional annotation file. By default, SAFE assumes that all node labels are systematic labels and thus `label_orf` can be populated directly from label. However, in three scenarios, SAFE populates `label_orf` from elsewhere:
  a. if the user is loading a network from a Cytoscape session file and the network has a node attribute named “ORF”;
  b. if the user is loading a network from a 5-column tab-delimited text file (see Note 1);
  c. if the default node labels in the network match gene names or allele names from *S. cerevisiae* and can be translated into their corresponding systematic ORF names.
6. As an alternative to the yeast GO biological process standard, the user can import any other annotation file that is provided in a tab-delimited matrix format. In this format, rows are node systematic labels, columns are functional attribute labels and the data are binary or quantitative values indicating the association between a node and an attribute. By default, SAFE will consider only the highest values of the attributes to be of interest: e.g., the 1s in a binary standard such as GO or the top positive values of a quantitative standard such as Hoepfner et al., 2014 (see Example 2). If only the lowest or both lowest and highest values of the attributes should be examined, the user should set `annotationsign` in `safe.ini` to `lowest` or `both`, respectively. Examples of binary and quantitative annotation files are provided as part of the SAFE package (`/data/` folder):
  a. `hoepfner_movva_2014_doxorubicin.txt`
  b. `hoepfner_movva_2014_hop_known.txt`
  c. `sample_annotation_file.txt`
7. Other distance metrics include `shortpath`, which measures the unweighted shortest path length between two nodes, and the `shortpath_weighted_edge`, which weights edges according to their original values supplied by the user. Unlike the default `shortpath_weighted_layout`, neither of these distance metrics depends on the layout algorithm and the resulting node positions.
8. Fisher’s exact test is used to calculate enrichments for binary functional annotations, such as GO terms. When the annotation standard is quantitative, enrichments are automatically calculated using random permutations and empirical p-values (see Example 2 for a more detailed explanation).
9. By default, neighborhood enrichment is assessed by comparing the frequency of an attribute within a neighborhood to the frequency of the attribute in the entire network (`background = map` in `safe.ini`). In case of functional biases in the network, enrichment can also be calculated relative to the total frequency of the attribute which is defined by the number of labels associated with the attribute in the annotation standard (`background = standard`).
10. In addition to `diameter`, the distance threshold can be defined in two other ways. The default `neighborhoodRadiusType` setting is `percentile` (with a default `neighborhoodRadius` value of `0.5`), which indicates that two nodes are part of the same neighborhood if their distance is in the lowest 0.5^th^ percentile of all pair-wise distances in the network. Alternatively, the user can define the distance threshold in absolute terms by setting `neighborhoodRadiusType` to `absolute` and `neighborhoodRadius` to a fixed value (e.g., 200). This option is particularly useful when node distances are measured by the unweighted shortest path length (nodeDistanceType = shortpath): for example, by setting `neighborhoodRadiusType` to `absolute` and `neighborhoodRadius` to `2`, the neighborhoods will contain only nodes that are within 2 hops away from each other.
11. The filter for region-specific attributes is set by `unimodalityType` in `safe.ini` and can have one of four values:
  a. `<empty>`: no filter is applied, all attributes are retained;
  b. `hartigans`: the Hartigan’s dip test of unimodality (Hartigan and Hartigan, 1985) retains a attribute only if the distribution of pair-wise distances between its significantly enriched neighborhoods is unimodal;
  c. `radius`: the radius-based method retains an attribute only if at least 65% of its significantly enriched neighborhoods are within a distance `2R`, where `R` is the distance threshold that defines the size of the neighborhood (see Step 7);
  d. `subclust`: the subtractive clustering-based method (Chiu, 1994; Yager and Filev, 1994) retains an attribute only if there is at most one cluster center among its significantly enriched neighborhoods.
12. Functional domains are groups of attributes that have similar enrichment landscapes. These groups are defined by a distance metric and a distance threshold (`groupDistanceType` and `groupDistanceThreshold` in `safe.ini`, respectively). The `groupDistanceType` can be set to `<empty>`, indicating that no grouping will be performed. Alternatively, it can be set to `jaccard` or `correlation` to compute distances via Jaccard similarity or Pearson correlation, respectively. Pearson correlation considers all enrichment scores, whereas Jaccard similarity takes into account only the significant ones. The `groupDistanceThreshold` defines the number and size of functional domains. It is expressed as a fraction of the total height of the hierarchical clustering tree and can assume any value between `0` and `1` (higher values indicate fewer and bigger clusters).
13. The domain filtering and consolidation step is optional and can be skipped by setting `groupMinimize` in `safe.ini` to `0`. Alternatively, two options are available: the default option (`groupMinimize = 1`) and an older “legacy” option (`groupMinimize 2`). The main difference between the two options is in how the predominant domain for a node is chosen. In the “legacy” implementation, domain X is predominant for a node if X is associated with the attribute that has the highest enrichment score in the node’s neighborhood. In the newer implementation, domain X is predominant for a node if X is associated with the largest number of attributes that are significantly enriched in the node’s neighborhood.
14. In this example, only the highest attribute values were considered of interest (see Note 6) and thus the output files are labeled with “highest” (e.g., `domain_properties_annotation-highest.txt`). If only the lowest values were considered, the files would be named “lowest”. If both lowest and highest values were included in the analysis, two sets of files would be printed.

## Conclusions and discussion

The rapid and systematic mapping of network domains enriched for specific functional attributes is a distinctive feature of SAFE that encourages deep exploration of biological networks through iterative annotation with multiple complementary functional standards. Here, I showed that the iterative annotation of the yeast genetic interaction similarity network with GO biological process and chemical genomic data can link genes and their functions to bioactive compounds and enable the prediction of novel mechanisms of drug sensitivity and resistance.

In contrast to other methods, SAFE was specifically designed to annotate complete networks rather than their separate parts (e.g., clusters). To achieve this goal, SAFE takes advantage of network layout algorithms that position nodes in two-or three-dimensional space based on an equilibrium of forces that reflects network topology. Using layouts, instead of clustering algorithms, is advantageous because they do not require hard boundaries between network domains and enable the annotation of sparse, yet functionally coherent, neighborhoods. A limitation of this strategy, however, is that our understanding of most layout algorithms is quite poor. It is yet to be determined which algorithm is the most appropriate for a particular network, which parameters produce the best results and how stable is the final outcome. Addressing these questions will unlock powerful comparative analyses of numerous biological networks and provide new insight into the global functional organization of the systems they represent.

## References

Baryshnikova, A. (2016). Systematic Functional Annotation and Visualization of Biological Networks. Cell Syst 2, 412-421.

Chiu, S. (1994). Fuzzy model identification based on cluster estimation. Journal of Intelligent and Fuzzy Systems 2.

Costanzo, M., Baryshnikova, A., Bellay, J., Kim, Y., Spear, E.D., Sevier, C.S., Ding, H., Koh, J.L., Toufighi, K., Mostafavi, S., et al. (2010). The genetic landscape of a cell. Science 327, 425–431.

Costanzo, M., VanderSluis, B., Koch, E.N., Baryshnikova, A., Pons, C., Tan, G., Wang, W., Usaj, M., Hanchard, J., Lee, S.D., et al. (2016). A global genetic interaction network maps a wiring diagram of cellular function. Science 353.

Cytoscape.org. Cytoscape User Manual. http://wiki.cytoscape.org/Cytoscape_3/UserManual-Cytoscape_3.2BAC8-UserManual.2BAC8-Navigation_Layout.Automatic_Layout_Algorithms (accessed on February 6, 2016)

DeRossi, C., Vacaru, A., Rafiq, R., Cinaroglu, A., Imrie, D., Nayar, S., Baryshnikova, A., Milev, M.P., Stanga, D., Kadakia, D., et al. (2016). trappc11 is required for protein glycosylation in zebrafish and humans. Mol Biol Cell 27, 1220–1234.

Giaever, G., Chu, A.M., Ni, L., Connelly, C., Riles, L., Veronneau, S., Dow, S., Lucau-Danila, A., Anderson, K., Andre, B., et al. (2002). Functional profiling of the Saccharomyces cerevisiae genome. Nature 418, 387–391.

Hartigan, J.A., and Hartigan, P.M. (1985). The Dip Test of Unimodality. The Annals of Statistics 13, 70–84.

Ho, C.H., Piotrowski, J., Dixon, S.J., Baryshnikova, A., Costanzo, M., and Boone, C. (2011). Combining functional genomics and chemical biology to identify targets of bioactive compounds. Curr Opin Chem Biol 15, 66–78.

Hoepfner, D., Helliwell, S.B., Sadlish, H., Schuierer, S., Filipuzzi, I., Brachat, S., Bhullar, B., Plikat, U., Abraham, Y., Altorfer, M., et al. (2014). High-resolution chemical dissection of a model eukaryote reveals targets, pathways and gene functions. Microbiological research 169, 107–120.

Kamada, T., and Kawai, S. (1989). An algorithm for drawing general undirected graphs. Information Processing Letters 31, 7–15.

Shannon, P., Markiel, A., Ozier, O., Baliga, N.S., Wang, J.T., Ramage, D., Amin, N., Schwikowski, B., and Ideker, T. (2003). Cytoscape: a software environment for integrated models of biomolecular interaction networks. Genome Res 13, 2498–2504.

Yager, R., and Filev, D. (1994). Generation of fuzzy rules by mountain clustering. Journal of Intelligent and Fuzzy Systems 2, 209–219.

